# Wearable Photonic Device for Multiple Biomarker Sampling and Detection Without Blood Draws

**DOI:** 10.1101/2025.01.23.634606

**Authors:** Zuan-Tao Lin, Biswabandhu Jana, Sandeep Korupolu, Yifei Kong, Guishi Liu, Yan Dong, Yongli Li, Quanwei Zhang, Wan Shou, Prabhat Upadhyay, Mei X. Wu

**Affiliations:** Wellman Center for Photomedicine, Massachusetts General Hospital, Department of Dermatology, Harvard Medical School, Boston, MA 02114, United States; Department of Mechanical Engineering, University of Arkansas, Fayetteville, AR 72701, USA

**Author notes:** Correspondence author: Dr. Mei X. Wu. ZL and BJ contributed equally to this work.

## Abstract

Blood draws or phlebotomy using needles have been practiced in clinics for several centuries and it often causes pain, discomfort, and inconvenience. Herein, we create a wearable photonic device by integrating a microlens array (MLA) and an optic microneedle array (OMNA) functionalized with immunobinding capacity for safe and needle-free biomarker sampling and detection. The MLA-integrated OMNA amplified and transmitted LED light at 595 nm into the skin through the OMNA, bypassing the light-absorbing melanin in the epidermal layer, and evenly distributing it in the capillary-enriched dermis. The 595 nm light was preferably absorbed by hemoglobin (Hb) and oxygen-Hb within the capillaries, triggering thermal dilation of capillaries without damaging them or causing petechiae. The light illumination resulted in remarkable increases in the concentrations of various blood biomarkers in the skin owing to capillary dilation and biomarker extravasation. These extravasated biomarkers bound specifically to the OMNA, which was covalently decorated with capture antibodies, each microneedle immobilized with one specific antibody. The multi-functional OMNA was extensively modified to amplify the immunobinding signals and achieve sensitivity superior to that of enzyme-linked immunosorbent assay (ELISA) kits. As a proof of concept, we validated the functionality of the prototype for minimally invasive sampling and precise multiplexed blood biomarker detection in two mouse models to quantify acute inflammation and specific antibody production. This cost-efficient device provides a promising platform for blood-free, multiplexed detection of blood biomarkers.

## 1. Introduction

Blood biomarkers are substances ranging from small molecules and ions to large entities like peptides, proteins, RNA, and DNA. These biomarkers are released into the circulation by cells, tissues, and organs during various biological activities within the body. Their concentrations in the blood circulation system reflect health and pathological statuses from specific tissues to the entire body.^[1]^ Therefore, the measurement of various blood biomarkers is prominent for disease diagnosis, prognosis, and monitoring.^[2]^ Current blood sampling, such as venipuncture, often causes pain and discomfort, especially for people with needle phobia or requiring frequent blood examinations.^[3]^ It can be done only by professional personnel and the efficiency highly relies on both the proficiency of a phlebotomist and the ease of accessing the patient’s vein. Moreover, healthcare practitioners who perform blood collection and testing face a risk of infections by bloodborne pathogens such as HIV.^[4]^ Furthermore, to date, blood tests are all carried out with expensive instruments and in well-equipped labs. Most tests are also time-consuming and require training lab staff, inconvenient for clinical practice, particularly in resource-limited regions. Therefore, there is a high demand and growing interest in point-of-care (PoC) tests that are low-cost, require minimal or no training, and need only small sample volumes via fingertip. In the past two decades, tremendous effort has been devoted to PoC test development, clinical testing, and commercial translation. Among them, the glucose meter, and pregnancy and Covid-test strips stand as the most successful ones.^[5]^ However, these tests measure primarily a single biomarker, greatly limiting their broad applications. Disease pathogenesis of many diseases often results from intricate biological processes that involve multiple signaling pathways. Thus, diagnosis, prognosis, and monitoring of disease onset and progression are not sufficient by a single biomarker. Multiple biomarkers are required for comprehensive insights to sufficiently assess pathogenic processes.^[2, 6]^ Clinical studies have demonstrated that multiple biomarkers can improve the preciseness of disease diagnosis considerably when multifactorial pathogenesis is taken into consideration.^[7]^

Microneedle array (MNA) comprising an array of microneedles with various geometries on a patch has attracted increasing interest in developing different biosensors or assays owing to its unique and promising properties including minimal invasiveness and sufficient surface area for functionalization.^[8]^ However, most current studies have developed MNA mainly for skin biomarker detection or sampling, rather than blood biomarker detection, making it primarily beneficial for skin diseases and sampling.^[9]^ is because the concentrations of blood biomarkers in the skin are either too low to be detected or significantly lower than in the blood. For instance, Wang et al. found that the IL-6 level detected by an MNA in mouse skin after injection of lipopolysaccharide (LPS) was 22-fold lower than that in serum.^[8c]^ Taking advantage of the preferable excitation of oxygenated hemoglobin (HbO_2_) and hemoglobin (Hb) in the wavelength range between 530 and 590 nm, physicians have utilized the lasers to selectively rupture blood vessels in human skin and treat symptoms of excess blood vessels for over two decades.^[10]^ Capillaries, composed of only a single layer of endothelial cells in their wall, are abundant in the upper dermis. We have shown that a laser with a very low energy level could induce controllable thermal dilation of capillaries allowing only blood biomarkers, not red blood cells, to extravasate and accumulate in the skin by 2,000 or 10,000-fold higher in the presence vs. the absence of laser treatment in mice and pigs, respectively.^[11]^ The photothermal induction of extravasation (PiE), followed by the insertion of a MNA functionalized with specific antibodies, enabled the uniform and reliable measurement of large molecules like antibodies or small molecules like FITC on each microneedle in the array,^[11a]^ raising an intriguing possibility that the MNA can potentially detect multiple biomarkers with one biomarker per microneedle in the array if a laser was used to pre-treat the skin. ^[11b, 12]^ In contrast, without laser treatment, only a few microneedles showed positive signals for biomarker binding, presumably resulting from uncharacterized leakage of biomarkers through capillaries damaged by penetrating microneedles as previously shown.^[11a, 13]^ Most microneedles in the array were negative or insufficient binding of biomarkers because concentrations of most blood biomarkers in the upper dermis are negligible or low. Such large variations among individual microneedles in the array make it impossible to quantify multiplex biomarkers in a single array. While increasing the length of the MNA could enhance the efficiency and reliability of blood biomarker measurement, deeper dermal penetration can cause pain and discomfort.

Although laser treatment of a tiny skin area for MNA insertion could greatly enhance the efficiency and reliability of blood biomarker sampling, skin color (mainly the melanin produced in the epidermis) and thickness significantly affect light penetration and the efficiency of biomarker extravasation. Apart from skin colors, light penetration of the skin is also influenced by different ethnic groups, body locations, and aging, all contributing to the variations in biomarker sampling through the skin using MNA. Moreover, the operation of a laser makes it difficult for home-use applications. Nowadays, light emitting diode (LED) has been established broadly in various medical applications due to the attractive features of safety, home uses, wearable devices, and cost-effectiveness. The non-coherent and non-monochromatic LED is the ideal alternative for low-level laser application.^[14]^ However, its limited penetration of the skin must be addressed to ensure that the light can penetrate the epidermis and reach the dermis without generating excessive heat, making it safe for home use. In this regard, Kim et al. aligned a microlens array and MNA and achieved a 9-fold improvement of light transmission in comparison with MNA for the treatment of skin infection.^[15]^

In this study, we designed and fabricated an MLA to confine light to the dermis through OMNA so as to minimize skin-associated variations and increase reliability of MNA-mediated sampling of blood biomarkers. Additionally, the surface of the OMNA was functionalized with signal magnification and specific capture antibodies in a one-antibody-per-needle fashion, creating an OMNA immunosensor. We also constructed a wearable and miniaturized photonic device aligned with an LED, MLA, and OMNA immunosensor to induce transient leakage of blood biomarkers beneath the skin. These extravasated biomarkers bound to the capture antibodies on the OMNA immunosensor for 20-30 min, after which the OMNA was removed from the skin and subjected to biotin-tyramide-mediated signal amplification assay (BT-MNA assay). To demonstrate its clinical potential, we validated the capability of this innovative platform in measuring C-reactive protein, inflammatory cytokine IL6, and specific antibody production induced by flu virus in two mouse models. To the best of our knowledge, this optical prototype represents the first example of multiple blood biomarker detection without blood drawing for home use.

## 2. Results and Discussions

### 2.1. Photothermal-induced extravasation device (PiED) and the underlying working principle

PiED was fabricated as a small, wearable device with a cylinder shape of 4 cm in diameter and 4.5 cm in height (Figure 1A and B). The prototype is designed to be flexible, with an adhesive backing for attachment to the skin (Figure 1A) and the photo is shown in Figure 1B. The LED light is aligned with the MLA and OMNA immunosensor within a chamber (Figure 1C). PiED can be separated into two major sections: “disposable MNA immunosensor” and “reusable photonics”. The cost-effective reusable photonics consists of all components except for an OMNA immunosensor. The MNA holder at the bottom of PiED makes it convenient to replace the OMNA immunosensor. Heatsink and micro-fan are used for cooling during LED illumination. As depicted in Figure 1D, mirroring the microneedle array, the microlens on MLA confine and direct light through each of the microneedles in the OMNA. Part of a light-up OMNA is shown in Figure 1E, in which the light was seen through and surrounding each microneedle. A temperature sensor is integrated into the PiED to turn off the light whenever the temperature rises to >43°C to ensure safety. PiED is equipped with a rechargeable battery via wireless charging. The illumination time can be pre-set.

**Figure. 1.**
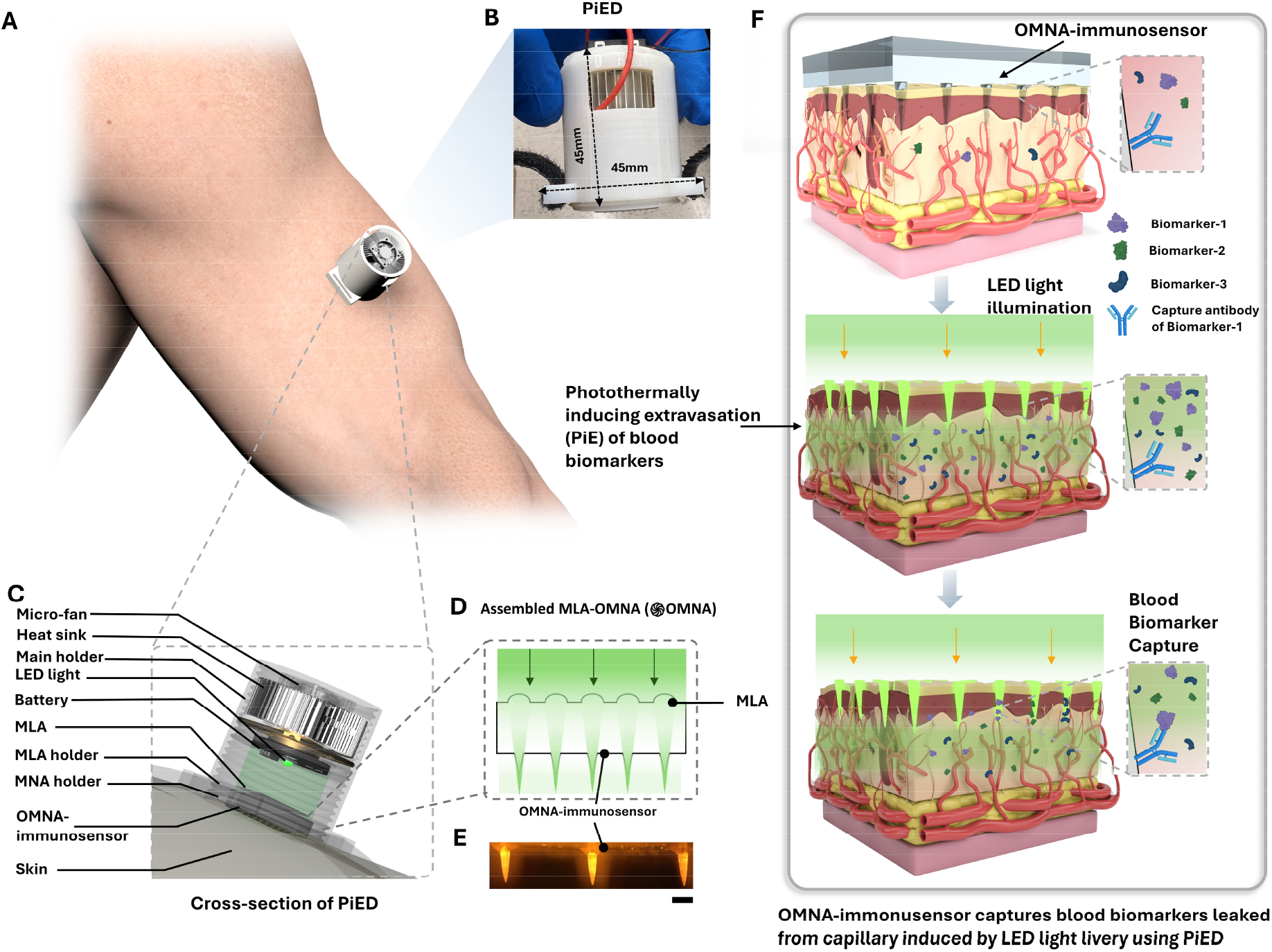
PiED for blood sampling and multiplex detection of blood biomarkers. **A**. Illustration of PiE on human arm showing its small size. **B**. The photo of the PiED showing its miniaturized size **C**. Cross-sectional illustration of the device with LED light illumination, and its major components. **D**. Illustration of light traveling through an assembled MLA-OMNA, allowing for light delivery evenly surrounding microneedles in deep skin tissue with capillaries. **E**. Side-view photographs of microneedles of the assembled MLA-OMNA (or ֍OMNA) under illumination of LED light. Scale bar: 100 μm. **F**. Schematics of mechanician of minimally invasive sampling through photothermally inducing capillary leakage of blood biomarker, which are then captured corresponding to capture antibodies on individual microneedle.

The concentrations of blood biomarkers in the upper dermis are too low to be detected by an OMNA immunosensor (upper, Figure 1F). However, following LED illumination (green light-up in individual microneedles), the light is delivered to the upper dermis and selectively excites HbO_2_ and Hb within the capillaries. This causes capillaries in the upper dermis to dilate, triggering the extravasation and accumulation of biomarkers only, but not blood cells (middle, Figure 1F). Thus, the levels of biomarkers increase dramatically in the upper dermis and bind to the specific capture antibodies immobilized on the surface of the OMNA immunosensor, with one biomarker per microneedle (bottom, Figure 1F).

### 2.2. Preparation of OMNA and its optical properties

Mechanical strength and optic properties are both required for an OMNA to insert the skin and deliver light through the epidermis. To achieve this, we employed polymethyl methacrylate (PMMA), a widely used transparent synthetic polymer known as “biocompatible glass”, to prepare an optic MNA (OMNA) by casting PMMA solution into a female polydimethylsiloxane (PDMS) MNA mold (Figure S1A, Supporting information).^[8b, 16]^ The resultant OMNA was transparent (Figure S1B and C, Supporting information). Its Young’s modulus is ∼3GPa, strong enough to penetrate the skin. In support, as shown in Figure 2E, the OMNA was readily inserted into the skin, as evidenced by a pattern of micro holes appearing in the skin only after, but not before, the insertion of the OMNA. Strikingly, after removal of the OMNA, the micro holes healed rapidly and completely by 30 min, with no bleeding or any petechiae observed, in agreement with capillary dilation rather than damage. This safe and minimally invasive biomarker sampling contrasts sharply with the microneedle-based blood collection device reported by Blicharz et al., which can lead to bleeding after device removal.^[3]^ The absence of any blood residue on the skin after the procedure is significant on several fronts: (1) No recovery time is needed; (2) the risk of infections for the patients is minimized; and (3) contamination of the environment or pathogen transmission is significantly limited.

**Figure. 2.**
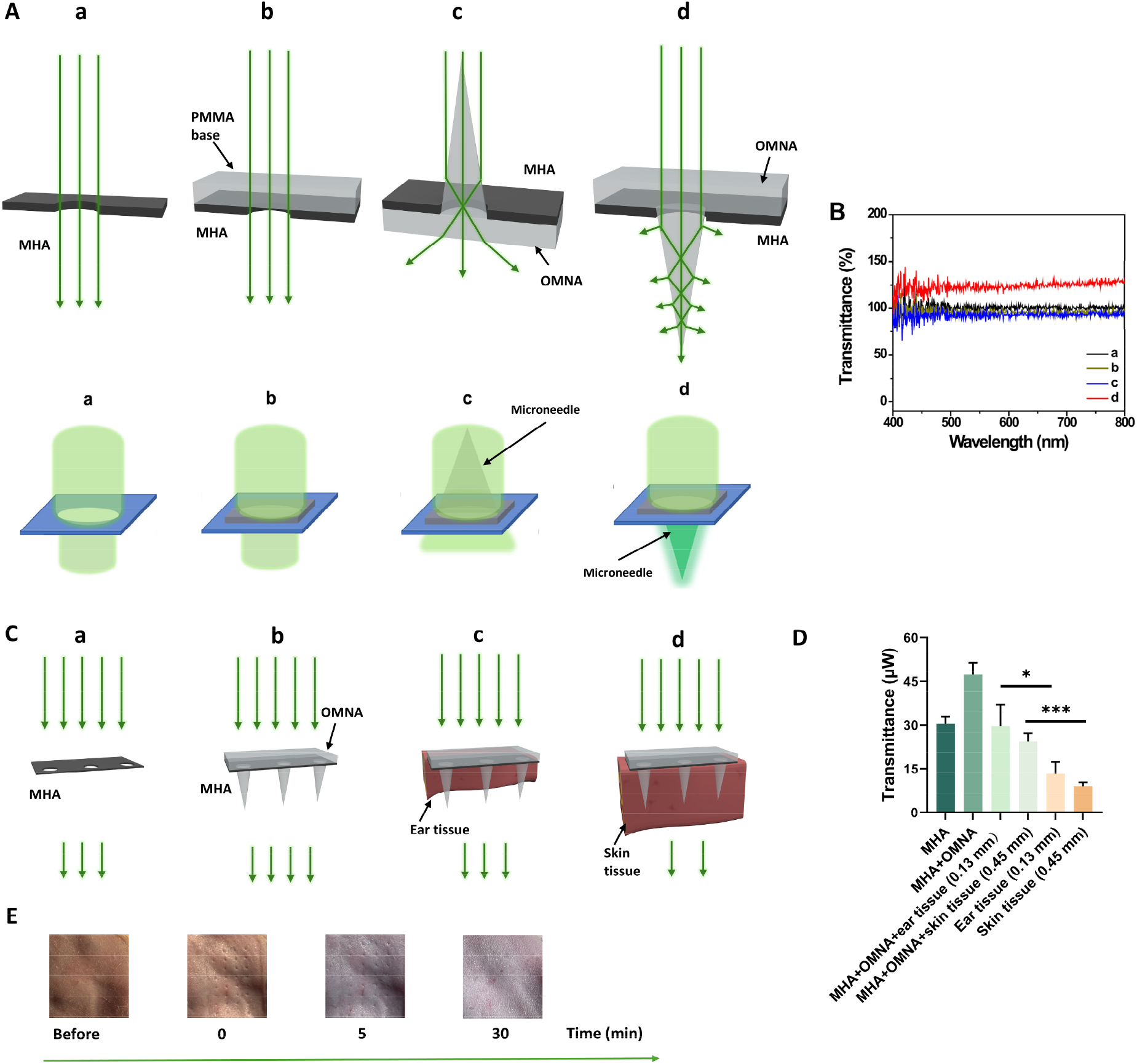
Optical properties and blood sampling of OMNA. **A**. Illustration of light travels through micro-hole array (MHA) made by aluminum foil **(a)**, PMMA **base (OMNA without microneedles) (b**), tip of the microneedles of OMNA and MHA **(c**), and bottom of the microneedles of OMNA and MHA **(d)**, and the possibility of corresponding path of light. **B**. The detected transmittance of the light with various wavelengths that goes through the different method in **A. C-D**. Illustration **(C)** and optical transmission **(D)** of OMNA on mouse ear tissue (0.13 mm) and mouse skin tissue (0.45 mm) using 25 mW LED light source. Data show mean ± s.d. **E**. Photographs of mouse skin before and after OMNA application at 0, 5, 30 min. Of note, no blood bleeding was observed. ^*^P = 0.0281, ^***^ = 0.0009, by two-tailed unpaired *t*-test.

To evaluate the light transmittance of the OMNA, we fabricated a micro-hole array (MHA) with aluminum foil and set the transmittance efficiency of light through MHA as 100% arbitrarily (Figure 2A a). The transmittance was slightly lower in the presence vs. the absence of the 1 mm thick PMMA base (without microneedles) (Figure 2A, b vs. a). However, against all expectations, the light transmittance rate was significantly increased through an OMNA (Figure 2A d and red in Figure 2B). An increase in the light transmission by the OMNA is presumably due to the cone-shaped microneedle that acts as a concave lens as illustrated in the upper panel, allowing the light travel through the microneedle at proper converging angles (Figure 2A d). The effect of the concave lens was proven because an inversion of the microneedle reduced the light penetration (Figure 2A c). The enhancement was demonstrated consistently over light spectra from the visible (400 – 700 nm) to near-infrared (700 – 800 nm) wavelengths (Figure 2B)

To assess the efficiency of light penetration of the skin through the OMNA, an OMNA was inserted into a 0.13 mm thick mouse ear tissue or a 0.45 mm thick mouse skin, followed by laser administering on top (Figure 2C). Amazingly, the laser transmission through the ear skin tissue was not compromised by the OMNA, bringing about the transmission efficiency comparable to or only slightly lower than that obtained in the absence of ear skin tissue or OMNA (Figure 2C a&c), which is attributed to the enhanced transmission by the OMNA (Figure 2C a&c vs b). Increased tissue thickness from 0.13 to 0.45 mm reduced laser transmission slightly but the laser transmission remains significantly high compared to mouse ear tissue or mouse skin without OMNA (Figure 2C c&d and D). Thus, the OMNA penetrates the skin and bypasses the epidermic layer directing the light to the upper dermis. This allows it to function independently of skin colors, locations, and aging, greatly improving the reliability.

### 2.3. Surface modification of OMNA to capture extravasated biomarkers in the skin

OMNA immunosensor was next developed by covalent conjugation of capture antibodies on the surface of the OMNA. Polyethylenimine (PEI) was utilized to coat the surface of PMMA after oxygen plasma treatment. The PEI-coating surface of OMNA was linked to different generations of dendrimers (G0, G4, G5, and G6) to further increase the functional groups for conjugation with abundant capture elements (Figure S2A, Supporting information). To determine the effect of different surface modifications, seven OMNAs with various surface modifications were fabricated and conjugated with the same amount of C-reactive protein (CRP) as a model biomarker. After the addition of a biotin-CRP-detection antibody and fluorescent dye-conjugated streptavidin, we measured the fluorescence intensity and found that the OMNA modified by G5 dendrimers had the best performance (Figure S2B, Supporting information). The OMNA immunosensor decorated with G5 dendrimers on PEI in each microneedle in the array was subsequently used in the studies (Figure S1B, Supporting information).

### 2.4. Design of MLA

While the OMNA could induce capillary dilation beneath the skin following 595 nm laser irradiation, the laser was relatively unsafe for home use. To replace the laser with a user-friendly LED, we must improve light transmittance to induce capillary dilation without generating excessive heat. To this end, we designed and fabricated a microlens array (MNA) mirroring the pattern of the OMNA. As depicted in Figure 3A, the MLA was placed on top of the OMNA, focusing and directing incoming light through the OMNA with one microlens per microneedle. The back focal length (*f*_*b*_) was calculated from the equation, ^[17]^

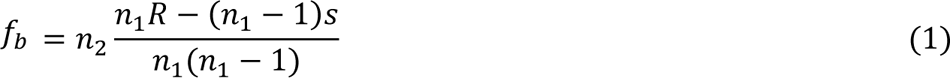

**Figure. 3.**
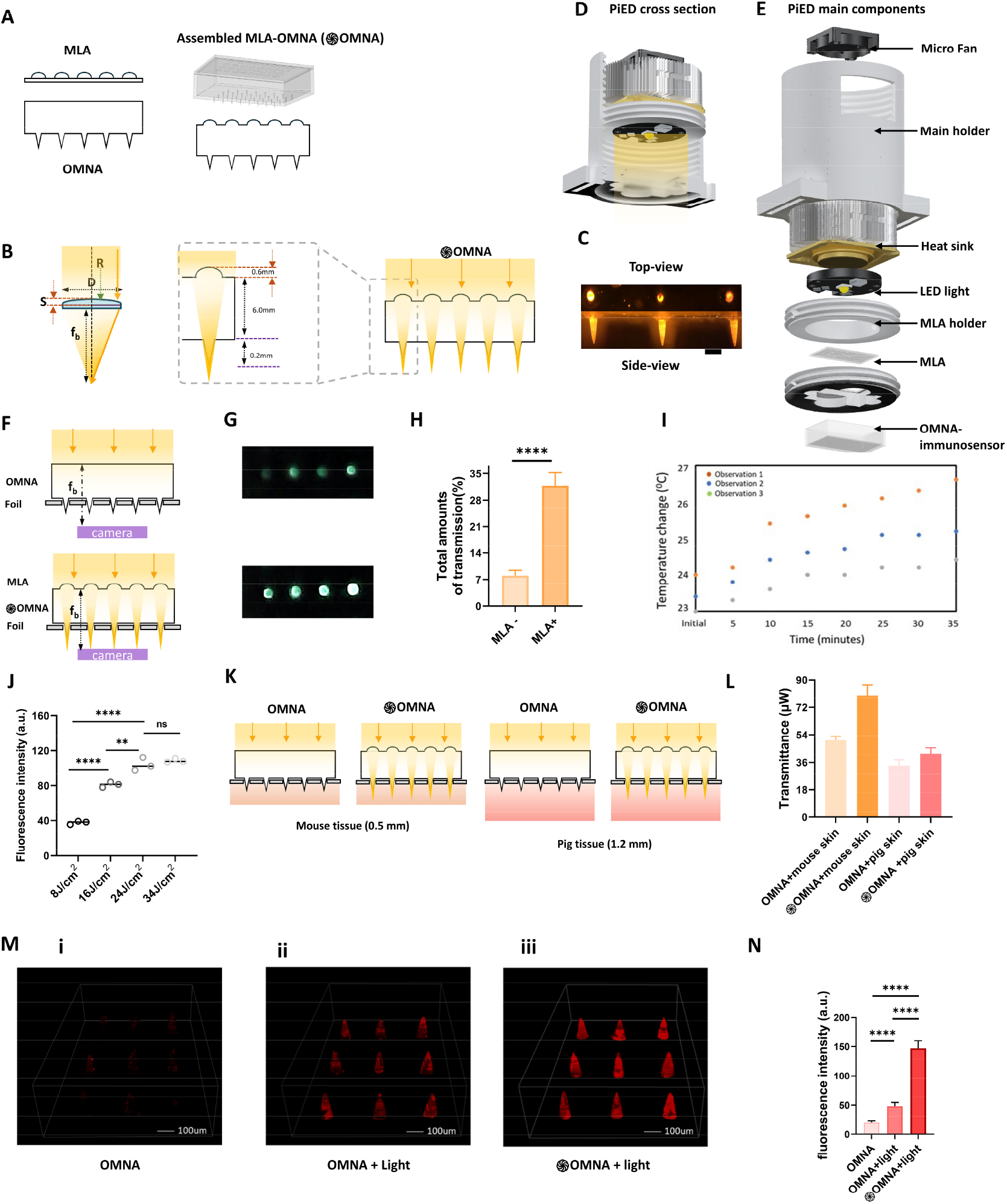
Components and optical characterizations of PiED. **A**. Side-view illustration of an assembled MLA-OMNA(֍OMNA), and the disassembled MLA and OMNA. **B**. Design of MLA. **C**. Cross-sectional illustration of the device with LED light illumination. **D**. Illustration of the components of the device. **E**. Side-view (top) and top-view (bottom) photographs of assembled MLA-OMNA under illumination of LED light. Scale bar: 100 μm **F**. Illustration of the experimental setup to measure light transmission through OMNA with (top) and without MLA (bottom) by using an aluminum foil mask punched by an array of holes aligned with microneedles. The foil mask prevents the background light transmission **G**. Optical images of light transmission with (top) and without MLA (bottom). **H**. Comparison of total amount of light transmission with and without MLA. **** P< 0.0001 by two-tailed unpaired *t*-test. **I**. Maximum temperature at single microneedle tip observed under illumination of a LED light source with a powder of 30 mV. **J**. Comparison of fluorescence intensity of microneedles of OMNA obtained from PiED applied on EB injected mice under illumination of LED light with different power levels after. ns, not significant. ^**^ P = 0.0074, ^***^ = 0.0001, ^****^ P< 0.0001 by two-tailed unpaired *t*-test. **K-L**. Illustration **(K)** and optical transmission **(L) of** PiED device with/without MLA on mouse tissue (0.5 mm) and pig tissue (1.2 mm) using 25mW LED light source. **M-N**. The EB injected mice were applied device with different experimental setup to explore the effect of MLA and light. Fluorescence images **(M)** and their corresponding intensity **(N)** of microneedles of from the OMNA of three PiED devices using different setups, including devices **(i)** containing only OMNA without MLA and light illumination, **(ii)** OMNA and light illumination, or **(iii)** (֍OMNA under illustration with a LED light at 24 J/cm^2. ****^ P< 0.0001 by two-tailed unpaired *t*-test. Data show mean ± s.d. NS, not significant. a.u., arbitrary units. Scale bar: 100 μm.

where 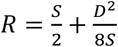 is the radius of curvature, S is the sag height, D is the diameter, n_1_ is the Lens material refractive index, and n_2_ is the air refractive index (Figure 3B). To ensure that light through the microlens and OMNA falls into the skin and obtains maximum light intensity surrounding the capillaries, targeting *f*_*b*_ should be slightly higher than 6.2 mm which covers the target distance (MLA base height of 1 mm + MNA base height + needle height) (Figure 3B). Accordingly, we fabricated MLA with a diameter of 1.5 mm, a sag of 0.1 mm, and a radius of curvature of 2.86 mm to obtain an effective focal length of 7 mm. The focal point is 1 mm and 0.8 mm lower than the surface of the skin, which is equivalent to the thickness of the human epidermal layer or the tip of the microneedle, respectively.^[15]^ The MLA was made via an MLA female mold created by 3D printing, to which PDMS and curving solution mixture was added, followed by heating for 4 h at 65°C (Figure S4A, Supporting information). After cooling down, MLA was obtained by peeling from the mold and assembled with the OMNA. We compared MLAs made from two different materials and found that PDMS MLA has the best performance. The 595 nm LED light source at 24 J/cm^2^ was delivered through microneedles in the assembled MLA-OMNA, as clearly seen from the side-view and top-view photograph of the OMNA immunosensor (Figure 3C).

### 2.5. Components and optical characterizations of PiED

The PiED was constructed by aligning three primary components: LED light, MLA, and OMNA immunosensor within the chamber of the main holder (Figure 3 D&E). The micro-fan and heat sink are used for cooling the LED. We validated the light transmission through an assembled MLA-OMNA (or ֍OMNA) for its effectiveness in the dilation of the capillaries. An aluminum foil mask was perforated with an array of micro holes aligning with individual microneedles to cover the OMNA base to exclude all light travelling through the base or outside the microneedles (Figure 3F). The intensity of transmitted light through ֍OMNA was significantly higher than that through OMNA without MLA (Figure 3G). OMNA increased the light transmission by 4-fold in the presence compared to the absence of MLA (Figure 3H). To assess the thermal effect of light transmission on ֍OMNA over time, a thermal camera was employed to measure the temperature of ֍OMNA in the presence of an aluminum foil mask at different time points. The temperature at the needle tip increased up to 3°C after exposure of 30 min, indicating only a slight heat generated by LED light (Figure 3I). To optimize the light fluence, we employed LED light sources with power at 8, 16, 24, and 32 J/cm^2^ to examine their effect on capillary leakage. The intensity increased proportionally with a rising light fluence up to 24J/ cm^2^, reaching a plateau thereafter (Figure 3J). Thus, 24J/ cm^2^ was identified as the optimized power source for assessing the optical transmission of the PiED device with/without MLA on 0.5 mm thick mouse tissue and 1.2 mm thick pig tissue (Figure 3 K-L). The light transmission of the device with ֍OMNA on mouse skin tissue showed the best performance, significantly higher than that of the OMNA alone. Although the light transmission decreased on pig skin tissue which is more than twice thicker than that of mouse tissue, the light transmission of the device with ֍OMNA was still higher than that of the OMNA alone.

To evaluate the biomarker extravasation, the fluorescence intensity of Evans blue (EB) on ֍OMNA was measured after insertion into the skin of an EB-injected mouse model. EB binds serum albumin in blood circulation following intravenous injection and is extensively used to detect vessel perfusion or damage of blood vessels by monitoring its leakage into the surrounding tissue. After binding with albumin, EB generates fluorescence in the red to far-red spectrum due to a conformational shift of a cis-trans isomerization.^[18]^ In brief, BALB/C mice were intravenously injected with EB and wore a PiED with a 30mW LED light source on lower dorsal skin after hair removal. The PiED was equipped with an OMNA without antibody conjugation. After turning on the PiED for 20 min, the EB-albumin complex was extravasated and could stick on the OMNA as imaged by a confocal microscope (Figure 3, M&N). The device setups included (i) containing only OMNA without MLA and light illumination, (ii) assembled MLA-OMNA without light illumination, or (iii) assembled MLA-OMNA under illustration with a LED light at 24 J/cm^2^. Notably, the PiED with ֍OMNA exhibited the highest fluorescence intensity with the light on. Compared to the device with only OMNA, the PiED-֍OMNA increased fluorescence intensity by 7.4-fold and 3-fold as compared to OMNA only or OMNA with light illumination, respectively (Figure 3N). The results indicate clearly that LED light successfully passed through ֍OMNA reaching the skin dermis resulting in the leakage of the EB-albumin complex.

### 2.6. Functionality of BT-MNA assay

We next established an MNA assay (BT-MNA assay) capable of quantifying the biomarkers captured on the individual microneedle in the OMNA immunosensor sensitively and reliably. First, the biotin-detection antibody mixture and streptavidin-horseradish peroxidase (HRP) were added onto the OMNA immunosensor, forming an HRP immunosandwich on each microneedle specifically (Figure 4A). Tyramide-biotin and hydrogen peroxide were then added, with the conversion of tyramide-biotin into an active product catalyzed by HRP and hydrogen peroxide, resulting in the localized deposition of biotin product at high density surrounding tyrosine residues on proteins or HRP (Figure 4A).^[19]^ This led to signal amplification when fluorescent dye-conjugated streptavidin was present. Biotin-tyramide-mediated signal amplification could substantially increase sensitivity and reduce the usage of antibodies and fabrication costs.

**Figure. 4.**
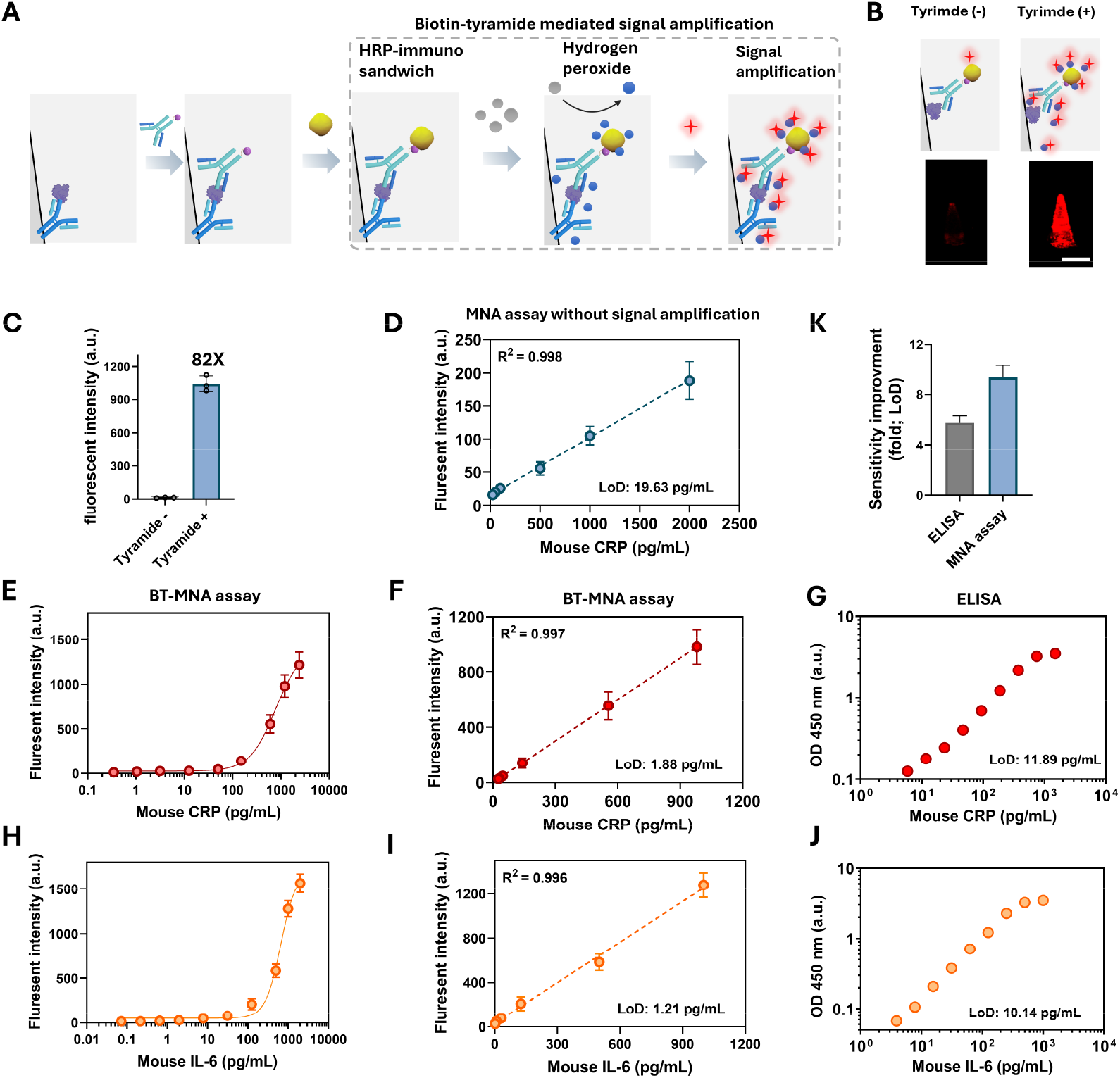
Functionality of BT-MNA assay based on OMNA-immunosensor. **A**. Illustration of biotin-tyramide mediated signal amplification. **B-C**. Illustration **(B, top)**, fluorescence images **(B, bottom)**, and corresponding intensity **(C)** of microneedles resulted from assay with **(B, right)** or without **(C, left)** biotin-tyramide mediated signal amplification. **D**. Standard curve for mouse CRP detection by MNA assay without signal amplification. Scale bar: 100 μm. **E-F**. Dose-dependent curve **(E)** and standard curve **(F)** of mouse CRP detection by BT-MNA assay. **G**. Standard curve for mouse CRP detection by ELISA kit. **H-I**. Dose-dependent curve **(H)** and standard curve **(I)** of mouse IL-6 detection by BT-MNA assay. **J**. Standard curve for mouse CRP detection by ELISA kit. **K**. the LoD BT-MNA is 5.7-fold and 9.4-fold lower than those of ELISA and MNA assay, respectively (without biotin-tyramide mediated signal amplification), Data show mean ± s.d. a.u., arbitrary units.

To verify the signal amplification by biotin-tyramide, we immobilized the same amount of CRP on two MNAs, followed by incubation of them with biotin-detection antibodies. One of them was assayed by streptavidin-HRP, biotin-tyramide, and hydrogen peroxide for signal amplification as above, while the other was assayed in the absence of these reagents used as control (Figure 4B top). The fluorescence intensity of individual microneedles revealed a much brighter fluorescent signal in the presence as compared to the absence of biotin-tyramide (Figure 4B bottom). As a result, an 82-fold increase in sensitivity was attained by the signal amplification mediated by biotin-tyramide (Figure 4C). To establish the ability of BT-MNA assay for sensitively detecting CRP and IL-6, two OMNA immunosensors were immobilized with capture antibodies directed at CRP or IL-6, respectively, for single biomarker detection, with the immunosensor assaying without biotin-tyramide amplifier as controls (Figure 4D). The standard curves of mouse CRP and IL-6 obtained by the BT-MNA assay offered the limits of detection of 1.86 pg/mL, and 1.21 pg/mL, and the linear ranges from 2.93 to 1500 pg/mL, and 1.95 to 1000 pg/mL, respectively (Figure 4F, I). In comparison to commercial ELISA kits, the BT-MNA assay increased the sensitivity by 6.4-fold, and 7.5-fold, respectively, for measuring CRP and IL-6. It is noteworthy that the linear ranges for CRP and IL-6 detection were much broader than those of ELISA. Table S1, Supporting information compares the functionality between BT-MNA assays and ELISA for quantifying CRP and IL-6. This increased sensitivity is attributed considerably to the biotin-tyramide-mediated signal amplifications. Without biotin-tyramide, the MNA assay was 9.4-fold less sensitivity (Figure 4K).

### 2.7. Preparation of an OMN-immunosensor for multiplex detection of biomarkers

To achieve multiple biomarker detection on one single OMNA immunosensor, a male mold was modified with a cylinder in replace of the microneedle geometry using a 3D printer to create a PDMS reaction female mold (Figure S3A, Supporting information). Each cylinder-shape hole in the PDMS reaction female mold functions as a micro-container and can hold a specific antibody/coupling reaction mixture solution of ? ul, facilitating the covalent conjugation of various capturing antibodies on individual microneedles in a one-specific capture antibody per microneedle fashion (Figure S3B, Supporting information).

### 2.8. Multiplex detection of biomarkers in mouse models

To demonstrate the clinical potential of the PiED, three biomarkers were quantified in two mouse models. Acute inflammation was modeled by intraperitoneal injection of lipopolysaccharide (LPS), while influenza A (H3N2) virus was intranasally administered. LPS, a primary compound found on the surface of most Gram-negative bacteria, leads to the generation of pro-inflammatory cytokines such as interleukin 6 (IL-6) in blood circulation by innate immune responses.^[20]^ Influenza virus can elicit antibody response specifically against hemagglutinin 3 (HA3), which can be measured by HA3 antigen immobilized on OMNA. On the other hand, both bacterial and viral infections stimulate inflammatory responses, resulting in the drastic elevation of CRP that serves as the critical biomarker for infections regardless of the nature of the pathogens.^[21]^ Therefore, three biomarkers including CRP, IL-6, and anti-HA3 antibody together could distinguish bacterial infections from flu viral infections as an initial diagnosis, which can greatly limit the antibiotic prescription, slowing down the development of multi-drug resistant bacteria.

To quantify the three biomarkers in a single OMNA, the OMNA immunosensor was simultaneously immobilized with HA3 antigen and capture antibodies specific for either CRP or IL-6 (Figure 5A). Then, we performed a cross-activity study to determine if there was any cross-reaction among the three selected biomarkers in the OMNA immunosensor. The assay was conducted with one targeted biomarker fixed at 25 pg/mL, while the other biomarkers with an increasing concentration from 0 to 400 pg/mL. The fluorescent intensity of the targeted biomarker had little change with elevating levels of the all co-existing biomarkers (Figure S6, Supporting information), suggesting little cross-activity in the assay. Since we need to apply the OMNA immunosensor on to skin of mice, OMNA immunosensor should detect the range of levels of CRP and IL-6 in mouse models. However, levels of CRP and IL-6 in mouse models are out of the linear range of BT-MNA assay mentioned above. Thus, we compromised the performance of BT-MNA assay to make to possible that all the measurements fall into the detectable range via adjustment of the amount of capture antibodies, which is shown in Figure S7, Supporting information. As illustrated in Figure 5B-C, we measured the biomarkers at different time points in the two mouse models with acute inflammation to be sampled after LPS injection, while specific antibody was measured two weeks after immunization. In the acute inflammation, we applied the PiED device onto the lower dorsal skin of mice with LED light on for 20 min for each sampling at 0h, 4h, 8h, and 24h after LPS injection. Blood was collected in the same group of mice at 8h and 24h. In the flu infection mice, we applied the device similarly on days 13 and 14 for sampling and also collected blood samples at the same time in these mice for comparison. The BT-MNA assays were conducted after removing the OMNA immunosensor from the device, while ELISA was carried out on serum samples as shown in Figure 5B-C. The location of the biomarker detection in the OMAN is depicted in Figure 5Di, and representative fluorescence images of the OMNA immunosensors are shown in Figure 5Eii-v.

**Figure. 5.**
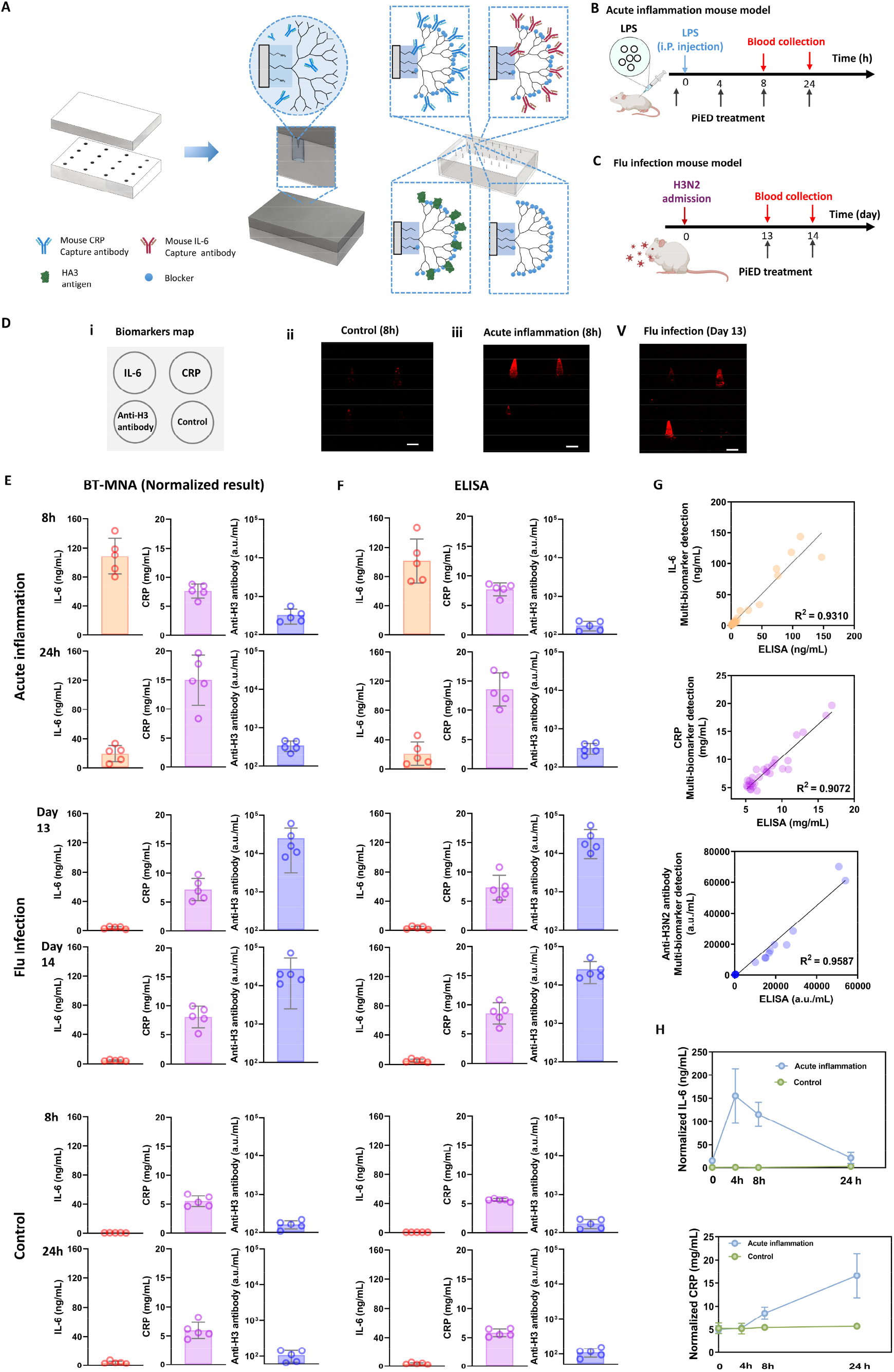
Multiplex detection of biomarkers in mouse models. **A**. Illustration of immobilization of capture elements (CRP capture antibody, IL-6 capture antibody, and HA3 antigen) on OMNA-immunosensor for detection of three selected biomarkers. **B-C**. Illustration of the workflow of immunization, blood sample collection, and application of PiED in two mouse models. (**B**)The mice were stimulated acute inflammation by i.p. injection with 2 μg/g LPS. We collected blood samples at 8h after LPS injection and on the second day at the end point of experiment. PiED was applied on mice at 0h, 4h, 8h, and 24h after LPS injection. (**C**) Mice were administered intranasally with flu H3N2 virus to induce flu infection. The blood sample collection and PiED treatment were carried out on Day 13 and Day 14 after administration of H3N2 virus. The mice injected with PBS was used for control. **D**. Biomarker map indicated the location of biomarkers on OMNA-immunosensor for multiple biomarker detection in mouse models (**i**), and the representative fluorescence images of OMNA-immunosensors obtained from PiED devices treated on different mouse models (**ii-v**). Scale bar: 100 μm. **E-F**. Levels of IL-6, CRP, or anti-H3 antibodies in serum measured by ELISA (**E**) and the corresponding normalized levels of IL-6, CRP, or anti-H3 antibodies which are multiply by dilution factors, measured by PiED in skin (**F**). **G**. The paired correlation test was carried out for the BT-MNA versus the ELISA results. **H**. The normalized levels of IL-6 and CRP in skin before and after capillary leakage were measured by PiED at different time points. Data show mean ± s.d. a.u., arbitrary units.

As shown in Figure 5E-F, we can find a significant higher level of IL-6 in the acute inflammation group than those in the control or flu infection groups. IL-6, the pro-inflammatory biomarker, increased dramatically in the first 8h after stimulation of acute inflammation, and reached a peak level at around 4h after administration of LPS, indicating the rapid response of the immune system of mice to defend infection (Figure 5H). Unlike human CRP, CRP levels were increased in both mouse models, but not dramatically. However, the level of CRP in acute inflammation group elevated significantly at 24h, which might be attributed to the induce of IL-6 (Figure 5H). Anti-influenza H3 antibody levels in the flu infection group were remarkably higher than those of the control and acute bacterial infection groups.

IL-6 was also measured by PiED device under different settings at 8h after LPS injection to examine the influence of components, and results agree with our findings mentioned above (Figure S8, supporting information). Without LED illumination and MLA or photo-induced extravasation, IL-6 in the skin was 5.012 ng/mL, while the serum IL-6 level determined by ELISA was 101.29 ng/mL, representative of 20.23-fold higher than that in the skin, in agreement with the findings of Wang et al.^[8c]^ The capillary leakage induced by LED light resulted in an 11.2-fold increase in IL-6 level detected by the MNA-immunosensor, reaching an average of 51.9% relative to the serum IL-6 level. Besides, the levels of CRP and anti-H3 antibodies in the skin were found to be 50% or 45% relative to the serum levels, respectively. Since the extravasated biomarkers were diluted by the skin interstitial fluid, we normalized the levels of biomarkers by multiplying the dilution factors, which were calculated in the basis of serum levels of the corresponding biomarker (Figure 5E). The dilution factors were estimated as 1.94, 1.99, and 2.21 for IL-6, CRP, and anti-H3N2 antibody, respectively. The normalized levels of biomarkers in the skin detected by PiED were well correlated with those in serum measured by ELISA in both mouse models (Figure 5G, E vs. F). The ability of the PiED-immunosensor to measure multiple blood biomarkers in a single application is of high clinical relevance. Unlike some diseases which can be monitored by a single biomarker, for instance, blood glucose levels for monitoring diabetes, most diseases’ pathological pathways are complicated and measurements of several biomarkers are essential for accurate and reliable diagnosis. The OMNA-based immunosensor provides an ideal approach not only in a minimally invasive manner but also for multiplex biomarker detection.

## 3. Conclusion

We have successfully designed and fabricated a wearable photonic device (PiED) by aligning LED light, an MLA, and an OMNA immunosensor. We characterized its optic properties and optimized its design to improve light transmission by MLA-mediated focusing and directing light to the capillary-enriched dermis through an OMNA, bypassing the epidermis and greatly diminishing the variations of light penetration of the skin owing to skin color, location, and aging. Importantly, instead of detecting skin biomarkers by reported MNA sensors, our device was optimized to trigger extravasation and accumulation of blood biomarkers, not red blood cells, beneath the skin. These extravasated biomarkers can bind directly on the OMNA immunosensor, offering an innovative platform to measure blood biomarkers without the need for blood draws. This technology enables safe and minimally invasive blood sampling and reduces the risk of exposure to infectious pathogens associated with conventional blood drawing. Moreover, by integrating an OMNA immunosensor, multiple biomarkers can be measured in a one-biomarker-one-microneedle fashion via BT-MNA assay, which can measure the selected biomarkers with higher sensitivity than conventional ELISA, enabling the detection of low-abundant biomarkers. The innovative platform can be easily adapted and expanded to detect dozens to hundreds of biomarkers for specific disease diagnosis and monitoring reliably and sensitively, holding great potential for multiple blood biomarker detection without drawing any blood.

## 4. Experimental Section

### 4.1 Materials

SYLGAR 184 Silicone Elastomer Kit containing polydimethylsiloxane silicone elastomers base and curing agent was purchased from Dow (Midland, MI, USA). Poly(methyl methacrylate) (PMMA) (Mw∼120000), poly(ethyleneimine) (PEI) (Mn∼60000; Mw∼750000), ethyl acetate, polyamidoamine (PAMAM) dendrimer (cystamine core, generations 2.0, 4.0, and 5.0), tris(2-carboxyethyl)phosphine hydrochloride (TCEP), 1-Ethyl-3-(3-dimethylaminopropyl)carbodiimide (EDC), N-hydroxysuccinimide (NHS), suberic acid bis(3-sulfo-N-hydroxysuccinimide ester), sodium hydroxide (NaOH), isopropanol, and biotin-tyramide were purchased from Sigma-Aldrich (Waltham, MA, USA). Streptavidin-horseradish peroxidase (HRP) was purchased from Abcam (Waltham, MA, USA), PBS from Life Technologies (Carlsbad, CA, USA), and MES buffer and Alexa Fluor 647-Streptavidin from the Thermo Scientific (Rockford, IL, USA). Mouse C-reactive protein (CRP) (DY1829), mouse IL-6 (DY406) DuoSet ELISA kits, and reagent diluent were purchased from R&D System (R&D Systems, Minneapolis, MN, USA). LED light (M595D3) was purchased from Thorlabs (Newton, NJ, USA), and norland Optical Adhesive 89 (NOA89) from (Jamesburg, NJ, USA). Distilled water was generated by using a Millipore Milli-Q ultrapure water purification system (Burlington, MA, USA).

### 4.2. Fabrication of optical microneedle array (OMNA)

An OMNA was made by modifying the method described previously.^[11a]^ In brief, PDMS elastomer base solution was mixed with curing agent (10:1) in one well of a 6-well plate. The plate was centrifuged at 2,000 rpm for 10 min to remove bubbles. MNA mold reported in our previous study was placed on the bottom of the well in the solution, followed by bubble removal under a vacuum and heating at 65°C for 4h. After cooling, the MNA mold was removed to obtain a female PDMS-MNA mold. PMMA was dissolved in ethyl acetate at 78°C and was stirred overnight to obtain a PMMA solution (20% w/v). The female PDMS-MNA mold was filled with 1 mL of PMMA solution and centrifuged at 4,000 rpm for 15 min, followed by heating at 85 °C until it was dried. The procedure was repeated 3 times, after which the OMNA was carefully peeled from the female PDMS-MNA mold.

### 4.3. OMNA surface modification

The resultant OMNA was washed with 2-propanol three times, air-dried, and treated by oxygen plasma using a Technics 500-II Plasma etcher (200 W) for 3 min, followed immediately by immersing in PEI solution (10% v/v, pH 11) with stirring for 6 hr. Next, PAMAM dendrons were prepared via cleaving 3 mM PAMAM dendrimer (cystamine core;) by 5 mM of TCEP. A desalting column was used to remove TCEP. The PEI coating on the surface of MNA was conjugated by PAMAM dendrons(2 µM) using suberic acid bis(3-sulfo-N-hydroxysuccinimide ester) sodium salt (2 µM).

### 4.4. Fabrication of optical microlens array (MLA)

In this work, we made MLA using PDMS or NOA89. An MLA female mold was designed in Solidworks and printed by a 3D printer. For fabrication of PDMS MLA, we fabricated MLA by PDMS elastomer base solution mixed with curing agent (10:1) using an MLA female mold. After heating at 65°C for 4h and cooling, the PDMS MLA was obtained by carefully separating from the MLA female mold. For fabrication of NOA89 MLA, NOA89 solution was poured on the mold and exposed using UV light at 365nm for around 90 sec. The ONA MLA was obtained by peeling from the MLA female mold.

### 4.5. Fabrication of PDMS reaction mold

A reaction male mold containing an array of tiny cylinders precisely aligned with the microneedles on MNA on a plain base was designed by Solidworks and printed by a 3D printer. PDMS base and curing agent were mixed well in a 6-well plate, to which the curing agent (10:1) was added. The plate was then heated at 65 °C for 4 h. The PDMS reaction female mold was obtained by removing the reaction male mold.

### 4.6. Fabrication of OMNA immunosensor by immobilization of capture elements on OMNA for multiple biomarker detection

After removing the reaction male mold, cylinder-shaped micro-reaction containers were created, and each of them could hold 2μL solution with one microneedle immersed within for immobilizing capture elements in one-capture element per microneedle fashion. IL-6 capture antibody, CRP capture antibody, or HA3 antigen was added into a mixture of EDC (0.4 M) and NHS (0.1M) in MES buffer (0.1M) and added into the designated micro-reaction container in the array. an OMNA was placed on top of the PDMS reaction mold to covalently conjugate IL-6 capture antibody, CRP capture antibody, or HA3 antigen on the surface of the designated microneedles in the OMNA through the EDC/NHS coupling reaction. After incubation with the mixture for 3h, the OMNA was removed and washed with a washing buffer made by PBS with 0.05% Tween-20 three times to remove unreacted reagents. The OMNA immunosensor was incubated with PBS solution with 2% skimmed milk (0.2 μm filtered) at 36°C for 1 h to block non-specific binding, followed by washing three times.

### 4.7. PiED fabrication

The main holder, MLA holder, and MNA holder were designed in Solidworks and printed by a 3D printer. LED light was connected to a heatsink with a micro-fan and battery and installed on top of the MNA chamber. The MLA and the OMNA immunosensor were attached to the top and the bottom of the MNA holder, respectively, assembling the OMNA with MLA for form the ֍OMNA. ֍OMNA were then installed in the bottom of chamber of main holder, allowing the alignment of LED light, MLA, and OMNA-immunosensor within the chamber of main holder.

### 4.8. In situ sampling and multiple biomarker detection in mice models

BALB/C mice at the age of 8 weeks were obtained from Jackson Laboratory. All animal experiments were reviewed and approved by the MGH Institutional Animal Care and Use Committee. The mice were randomly divided into three groups: Acute inflammation group, flu infection group, and control group. To stimulate acute inflammation, mice were intraperitoneal injected with 2 μg/g of LPS. To model flu infection, mice were Intranasally administrated of 10x LD50 of influenza A Aichi H3N2. As a control, mice were intraperitoneal injected with PBS. Prior to treatment of PiED, the hair was removed about 1-2 cm^2^ of the lower dorsal skin of the mice.

Subsequently, PiED devices were applied on mice with acute inflammation before LPS injection, and 4h, 8h, and 24h after injection by LPS. The PiED device containing a 595 nm LED light source at 24 J/cm^2^ was fastened on the dorsal skin for 20 min. Once the PiED device was removed, facial vein blood collection was immediately conducted at 8h and at 24h after injection by LPS. The OMNA-immunosensor was then detached from the device and stored at 4°C, and the serum obtained from blood samples were stored at -80°C for further investigation.

For flu infection group, the mice were treated by PiED devices on day 13 and day 14 after influenza H3N2 infection. The blood samples were immediately collected by facial vein after application of mMLA-OMNA was done.

### 4.9. BT-MNA assay for multiple biomarker detection

Different concentrations of mouse IL-6, mouse CRP, or bovine serum albumin (BSA) in 2% BSA in PBS were added onto the OMNA immunosensor in the micro-reaction containers described above. After a 2h incubation at room temperature (RT), the OMNA immunosensor was rinsed with washing buffer three times to remove the non-specific binding, followed by adding the biotinylated detection antibodies and incubation for 1h in the micro-reaction containers. After washing three times, the streptavidin-HRP was added and incubated for 20 min at RT, followed by three washes. The OMNA immunosensor was further incubated with 6 μg/mL of biotin-tyramide in 0.003% hydrogen peroxide in borate buffer for 30 min at RT. This was followed by 30-minute incubation with streptavidin-Alex Fluo 648 at RT in a dark room. After three washes, the fluorescence images of the OMNA immunosensor were captured by an intravital two-photon confocal microscopy (Olympus FV-1000).

For multiple biomarkers detection in mouse models, the OMNA immunosensor removed from the PiED was incubated with a mixture of biotinylated detection antibodies directed at mouse IL-6 or CRP and biotinylated anti-mouse IgG for 2h at RT. The following steps were the same as described above.

### 5.0. Statistical analysis

Two two-tailed t-tests were employed to evaluate the statistical difference between two groups. One way ANOVA was utilized to analyze the statistical difference among multiple groups. P value was determined and figures were plotted by GraphPad Prism 9 (GraphPad, CA, USA). The LOD was obtained by calculating the level of biomarker by using the sum of the mean fluorescence intensity of blank and 3 σ. All values are expressed as mean ± standard derivation (s.d).

## Acknowledgment

This work was supported, in part, by the Defense/Air Force Office of Scientific Research, Military Medical Photonics Program under award number FA9550-23-1-0656 and FA9550-20-1-0063 and Department discretionary funds to M.X. W. We are grateful to Calixto Saenz and The Microfluidics Core Facility at Harvard Medical School for their support in OMNA surface modification.

## Data Availability

All data supporting the results in this study are displayed within the paper and the Supplementary Information.

**Supporting Information Supporting information** is available from the Wiley Online Library.

