## Supplementary Information for "Wearable Photonic Device for Multiple Biomarker Sampling and Detection Without Blood Draws"

* Correspondence author:

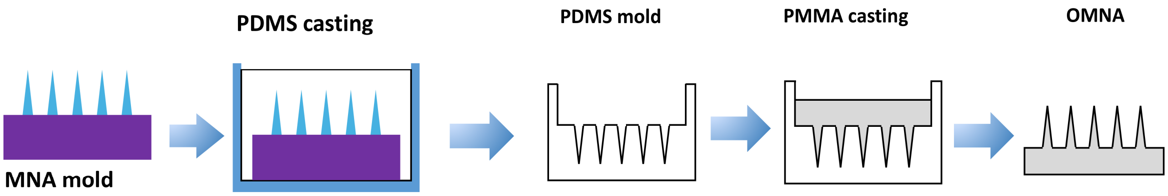


**Supporting information Figure S1. A.** Illustration of OMNA fabrication**.**

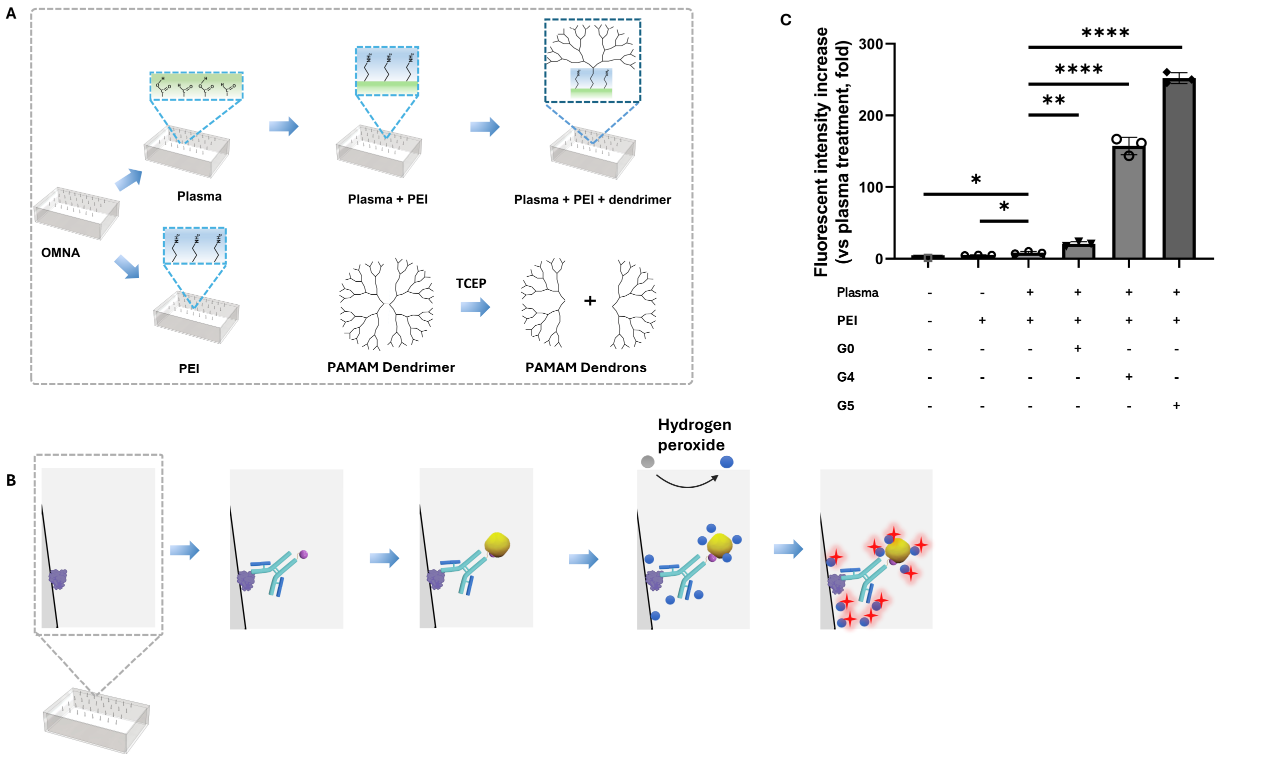


**Supporting information Figure S2. A.** Illustration of OMNA with different surface modifications, and the preparation process of PAMAM dendrons from PAMAM dendrimer. **B**. Workflow developed to evaluate fluorescence intensity. **C**. Comparison of fluorescence intensity generated by OMNA with different surface modification. Data show mean ± s.d. NS, not significant. * P = 0.0145, ** = 0.0037, **** P< 0.0001 by two-tailed unpaired *t*-test. a.u., arbitrary units.

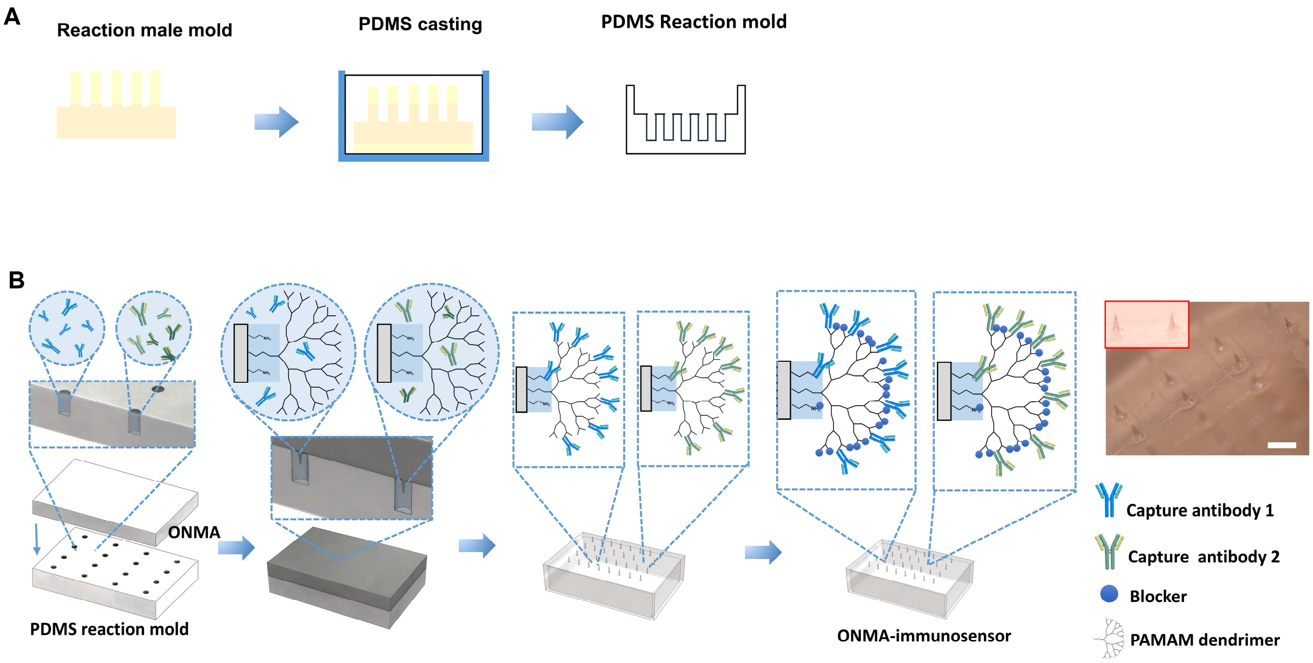


**Supporting information Figure S3. A**. Illustration of fabrication of PDMS micro-reaction container. **B**. The strategy of capture elements immobilization on OMNA in one-capture element-one-microneedle fashion. The hole resulted from the 3D printed reaction male mold, which is precisely aligned with microneedle on OMNA, creates a micro-reaction container for filling with mixture of specific capture element/coupling reaction for conjugating specific capture element on specific microneedle on OMNA.

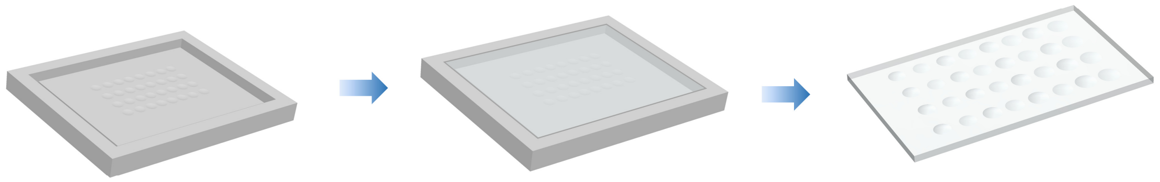

**Supporting information Figure S4.** Illustration of MLA fabrication.

.

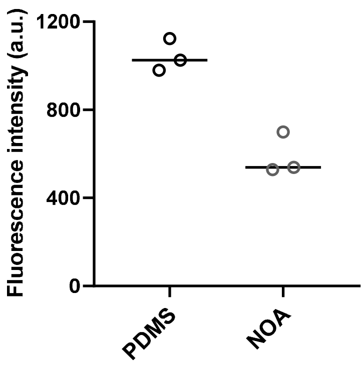


**Supporting information Figure S5.** Comparison of fluorescence intensity of microneedles of OMNA obtained from PiED applied on EB injected mice with different materials under illumination of LED light at 24 J/cm$^2$.

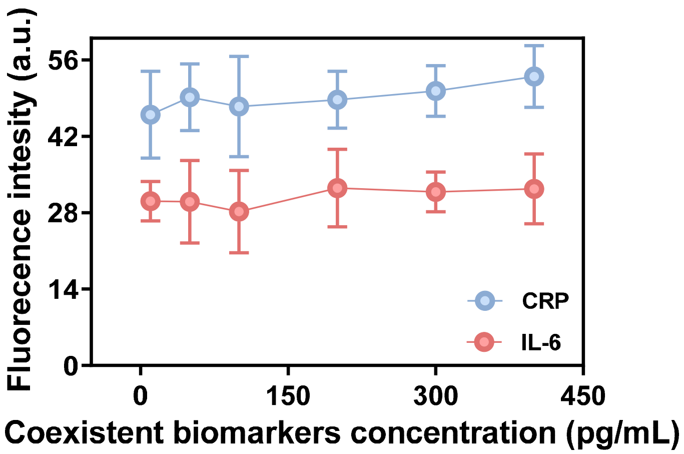


**Supporting information Figure S6.** Cross-reactivity study. We used the same level of detected biomarker at 25 pg/mL, and elevated the level of other biomarkers from 10 to 400 pg/mL to obtain various solutions for testing the OMNA-immunosensor.

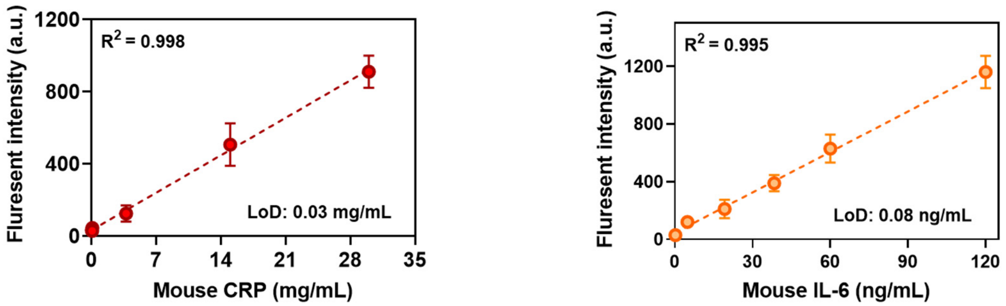


**Supporting information Figure S7.** Standard curve for detection of mouse CRP and IL-6 used in BT-MNA assay for multiple biomarker detection.

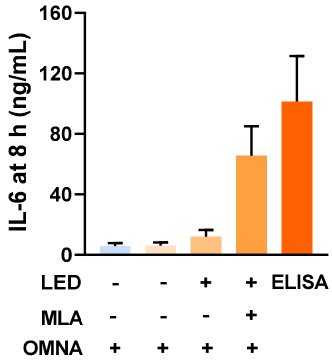


**Supporting information Figure S8.** Comparison of levels of mouse IL-6 detected by PiED devices with different experiment setups and ELISA kit.

**Supporting information Table S1**. Comparison of functionality of BT-MNA for detection of mouse CRP and IL-6 with that of corresponding commercial ELISA kits

| Assay | LOD | Linear Range |
|---|---|---|
| Mouse CRP ELISA Kit (R&D) | 11.89 pg/mL | 23.4 - 1,500 pg/mL |
| **Our work** | **1,88 pg/mL** | **2.93 - 1500 pg/mL** |
| Mouse IL-6 ELISA kit (R&D) | 10.14 pg/mL | 15.6 - 1000 pg/mL |
| **Our work** | **1.21 pg/mL** | **1.95-1000 pg/mL** |
